# Early-life social instability affects aggression, but not response inhibition, in chickens

**DOI:** 10.1101/2025.10.29.685273

**Authors:** Kathryn Willcox, Alizée Vernouillet, Birgit Szabo, An Martel, Luc Lens, Frederick Verbruggen

## Abstract

Living in complex social environments presents behavioural and cognitive challenges. A key challenge is the regulation of aggression, which may be facilitated by response inhibition, the ability to suppress inappropriate actions. In more complex social environments, greater variability in interactions may increase demands for aggression regulation, promoting the development of response inhibition. However, social instability, a form of complexity characterised by frequent changes in group composition, can elevate aggression and may instead impair inhibition. To test these competing predictions we raised 144 chickens (*Gallus gallus domesticus*) in either stable or unstable social groups for five weeks. We observed the aggression within their enclosures following the final manipulation, and assessed their response inhibition using a thwarting task and a cylinder task. Unstable groups initially showed more aggression than stable groups following regrouping, but this declined rapidly to the same lower level of aggression as the stable groups. Contrary to our predictions, there were no differences between birds raised in stable and unstable groups for any measure of response inhibition, and individual variation in aggression did not correlate with inhibitory performance. These findings suggest that social instability influences aggression but does not affect the development of response inhibition in chickens, highlighting the potential context- and species-specific nature of social effects on cognition.

## Introduction

Group living presents behavioural and cognitive challenges, as individuals must navigate the demands of cooperation and competition with conspecifics (Byrne and Bates 2007; Wascher et al. 2018). These challenges may differ depending on the nature of the social environment. Social systems vary widely both within and across species, and those in which individuals encounter a greater number and diversity of social interactions are typically considered to be more “complex” due to their variability (Kappeler 2019; Freeberg 2012; Rebout et al. 2021). Such social environments are thought to impose greater demands on individuals, potentially requiring more flexible behaviour and enhanced cognitive capacities to cope with increased social challenges (Byrne and Whitten 1988; Shultz and Dunbar 2007; Byrne and Bates 2007; Taborsky and Oliveira 2012; Amici et al. 2008; Wascher et al. 2018).

Aggression is a key challenge in social groups. Aggressive interactions between group members can be functional, granting individuals access to resources (e.g. Overduin-de Vries et al. 2020; Shimmura et al. 2007; Curren et al. 2015). In some species, aggression is also involved in the formation and maintenance of dominance hierarchies, which provide social structure, reduce further intense conflict and promote cooperation (Tibbetts et al. 2022; Holekamp and Strauss 2016; Rittschof and Grozinger 2021). However, aggression can also be costly, potentially causing injury or stress. Therefore, group-living animals must regulate when to behave aggressively (Smith and Price 1973; Clutton-Brock and Parler 1995; Asahina 2017; Estevez et al. 2007; Gersick et al. 2012; Wright et al. 2019). The demands for such flexibility in aggressive behaviour are likely to be heightened in more variable social environments (Gersick et al. 2012; Sachser et al. 2018; Loggia and Taborsky 2024).

A potential cognitive control mechanism involved in the regulation of aggression is response inhibition, the ability to inhibit inappropriate prepotent behavioural responses. In humans, deficits in response inhibition have been associated with heightened aggression (e.g. Madole et al. 2019; Puiu et al. 2018). Similarly, in non-human animals, individuals that are more aggressive have been found to have a lower ability to inhibit their actions across several species (e.g. Overduin-de Vries et al. 2023, Gobbo and Semrov et al. 2022; Rudebeck et al. 2007; Cervantes and Deville, 2007; Vernouillet et al. 2025). If response inhibition is involved in regulating aggression, we could, therefore, expect it to be enhanced in more complex social environments, where individuals must flexibly manage aggression under greater social variability. There is evidence for this: at an evolutionary scale, species with more complex social systems tend to be better at inhibiting actions than related species with simpler systems (Amici et al. 2008; Amici et al. 2018; Joly et al. 2017; Loyant et al. 2023). Similarly, at a developmental level, individuals exposed to greater social complexity often perform better on inhibition tasks than those from less complex environments (Ashton et al. 2018; Johnson-Ulrich and Holekamp 2020; Vernouillet et al. 2025), supporting the idea that social complexity may drive differences in response inhibition.

The relationship between social complexity and response inhibition is, however, not consistent across studies, with results varying across species, inhibition tasks, and measures of social complexity (Vernouillet et al. 2025; Willcox et al. 2024; Lucon-Xiccato et al. 2022; Triki et al. 2024; MacLean et al. 2014). This suggests that the link between the social environment and response inhibition is context-dependent. Specifically, different aspects of social complexity may place distinct demands on aggression regulation. For example, in larger groups (often considered more socially complex), several species show reduced aggression, potentially switching to a socially tolerant strategy (e.g. Syarifuddin and Kramer, 1996; Færevik et al. 2007; Estevez et al. 2007; Estevez 1997). Conversely, *unstable* social environments, in which group composition frequently changes, can increase aggression (e.g. Arnould et al. 2024; Koert et al. 2021; Andersen et al. 2008; Väisänen et al. 2004). In such competitive and unpredictable social contexts, a lack of inhibition could be an adaptive strategy to gain access to resources (Stevens and Stephens 2010; Fenneman et al. 2022; Eben et al. 2024). Supporting this idea, the only study to date that experimentally manipulated early-life social stability found that guppies (*Poecilia reticulata*) raised in stable groups showed better response inhibition than those from unstable groups (Lucon-Xiccato et al., 2022). However, fission–fusion social systems represent a natural form of social instability that can instead favour greater response inhibition, as individuals must adjust their behaviour depending on the individuals currently present (Amici et al., 2008; Amici et al., 2018; Aureli et al., 2008). Thus, the influence of social instability on response inhibition likely depends on species’ social systems and the specific behavioural demands they impose.

Here, we examine how early-life social instability affects both aggression and response inhibition, to further understand how this aspect of social complexity influences both behaviour and cognition. We used slow-growing broiler chickens (*Gallus gallus domesticus*), an ideal model for such research, as many individuals can easily be raised simultaneously without adults in controlled conditions. Chickens are social animals; in small groups they develop dominance hierarchies known as pecking orders (Guhl 1958), in which dominant individuals have priority access to resources (Shimmura et al. 2007). Thus, chickens have a social system in which response inhibition is likely to be necessary (Johnson-Ulrich and Holekamp 2020). To manipulate social stability, chicks were raised in either stable groups (same individuals kept together) or in unstable groups (group membership changing regularly). At the end of the manipulation, we recorded the birds’ aggressive behaviour within their enclosures.

The conflicting findings of previous studies could be due to the use of different tasks and measures to assess response inhibition. Therefore, we used two tasks to evaluate the effect of social stability on response inhibition in chickens. First, we used a ‘thwarting’ task (Troisi et al. 2025; similar to that used by Lucon-Xiccato et al. 2022) in which food is visible but inaccessible; there is no solution to obtain the reward and the animal is required to completely cease an ineffective behaviour (i.e. foraging attempts) in one trial. A shorter time spent persisting in trying to reach the food is considered better response inhibition, as the animal has been able to rapidly inhibit repeating an ineffective action. Second, we used the cylinder task, which has previously been used in studies exploring the influence of social environments on response inhibition (MacLean et al. 2014; Ashton et al. 2018; Johnson-Ulrich and Holekamp 2020; Willcox et al. 2024; Vernouillet et al. 2025; Triki et al. 2024; Guadango and Triki 2024). Unlike the thwarting task, the cylinder task has a solution; the animal is required to inhibit a prepotent behaviour (pecking the cylinder) to change to an effective behaviour (reaching through the open sides). Previous studies have shown chicken and red junglefowls’ (*Gallus gallus*) performances on the cylinder task to improve over multiple trials (Ferreira et al. 2020; Ryding et al. 2021), showing that they are able to learn to inhibit their actions in this task. Thus, we used this task to also assess whether such learning to inhibit is affected by early-life social stability.

We expected to observe more aggression in unstable groups than in stable groups because chickens (including broiler strains) display heightened aggression when interacting with unfamiliar individuals (Väisänen et al. 2004; Arnould et al. 2024). Given the proposed link between response inhibition and the regulation of aggressive behaviour, we predicted that early-life social instability would impair response inhibition. Specifically, we expected chicks from unstable groups to spend more time pecking during the thwarting and the cylinder tasks, and to be slower to learn during the cylinder task, compared to chicks from stable groups. Finally, we conducted an exploratory analysis to examine whether individual differences in aggression were related to variation in response inhibition, to provide further insight into the role of response inhibition in regulating aggressive behaviour within different social environments.

## Methods

### Subjects and Housing

A total of 144 slow-growing broiler chickens (Sasso XL451) were used for this study. This strain was chosen as they are large enough to carry ultra wide-band trackers (Salas et al. 2024) at 6 weeks old (required later as part of a related study), but are a slow-growing strain that has higher activity levels than standard commercial broilers (Stadig et al. 2018; Tickle et al. 2018), minimising any restricting influence of rapid growth on their behaviour.

The birds were raised in four batches of 36 chicks – two batches experienced the stable social treatment and two experienced the unstable social treatment. Data was collected in July - October in both 2023 and 2024, with one stable batch and one unstable batch raised each year. Within each year, the raising of the two batches was staggered to make testing feasible, and the order of the two batches was counterbalanced across the two years.

For each batch, 40 one-day-old sexed chicks were obtained from the commercial hatchery L’Oeuf d’or (Ardenne, Belgium) and brought to ECoBird’s research facilities at the Wildlife Rescue Centre Ostend (Belgium) to be raised. Upon arrival, the chicks were randomly allocated to groups of ten with an even sex ratio. Each batch was housed across two indoor lofts - each loft was further divided in two with wooden panels and an opaque mesh, to create a total of four enclosures of 2m x 1.5m.

For the first week (day 1-7; Fig. 1), each group of ten was housed in a smaller cage (91 x 53 cm) placed within one of the four enclosures. The cage contained two electric hen heating plates (Comfort Chicks), a water dispenser (Aveve), and a chick feeding trough (Voss Farming). Food (Wielink Kuikenmeel, Junai) and water were provided ad lib. The birds were handled every other day to habituate them to handling, and dried mealworms were gradually introduced. This acclimatisation week was included to avoid mortality and allow time for the chicks to become familiar to one another.

**Fig. 1.**
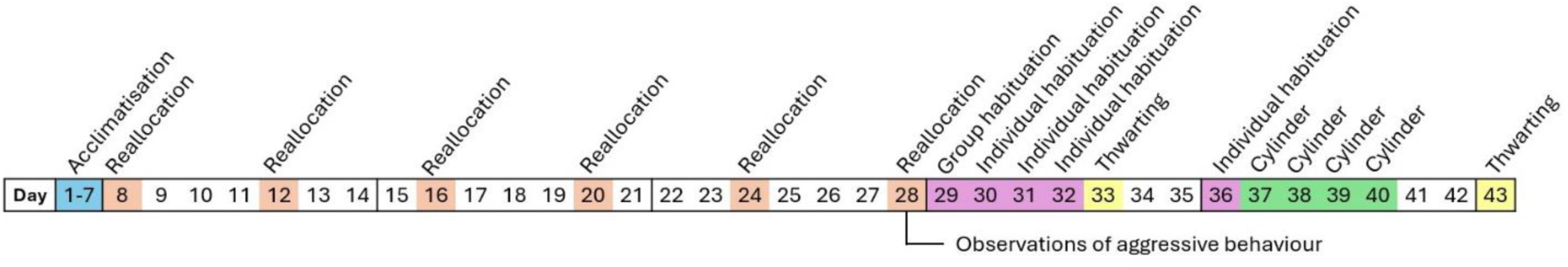
Timeline of the study, including reallocations, behavioural observations, habituation and testing.

After the first week (day 8 onwards; fig. 1), the birds were housed in the full size enclosures. At this stage, one bird from each group of ten was removed at random to form the experimental groups of 9. Our target group size was eight, but groups were initially established with ten chicks to allow for potential early-life mortality. As mortality was lower than anticipated, we reduced each group to nine birds at the start of the manipulation (by randomly removing one chick) to align with the planned movement of birds in smaller subgroups. Surplus chicks were kept for 4 more days as potential replacements to maintain consistent group sizes in case of mortality; one such replacement was made in the unstable treatment group in 2023 (day 12). Each enclosure contained wood-chip substrate and the following materials: a circular feed silo (⌀ 30cm, Junai), a sand bath, two electric hen heating plates (Comfort Chicks), a perch with two levels (50cm wide, one perch 20cm high, one 40cm high), and multiple water dispensers (Junai) providing water ad-lib throughout. Extra feeders were added to each enclosure to accommodate the growing birds on day 29 (after the social stability manipulation, Fig. 1).

Coloured leg rings were placed on the chicks’ legs to identify individuals; these were replaced as necessary as the chicks grew. The birds were visually checked multiple times a day, and could also be viewed remotely via cameras (Bascom) placed above the enclosures. Ambient temperature and humidity for each loft was monitored and recorded daily, to ensure they remained within those suitable for the age of the birds (Commission of the European Communities, 2007, document 2007/526/EC). During days 1-4 the lights were on from 6am-9pm; from day 5 onwards they were on from 7am-7pm (12hrs); there was no natural light. The birds’ weight and tarsus length were measured on days 8, 28 and 44.

### Manipulating Social Stability

The social stability manipulation began on day 8 and continued until day 28 (Fig. 1). Every four days, all birds were reallocated to a new enclosure. For the stable batches, the birds were always placed with their original social groups. For the unstable batches, the birds were placed in new combinations of nine birds each time (Fig. 2). The location of each bird after each reallocation was pre-determined; for the unstable groups, the predetermined reallocations were calculated to minimise the frequency of repeat encounters as much as possible (via the ‘Good Enough Golfers’ tool - Buchanan 2017), restricting any two birds from meeting more than three times. The nine birds within an enclosure each had a different coloured sticker placed on their back, so that individuals could be identified via the cameras placed above the enclosures. A full description of the reallocation procedure can be found in the Supplementary Information.

**Fig. 2.**
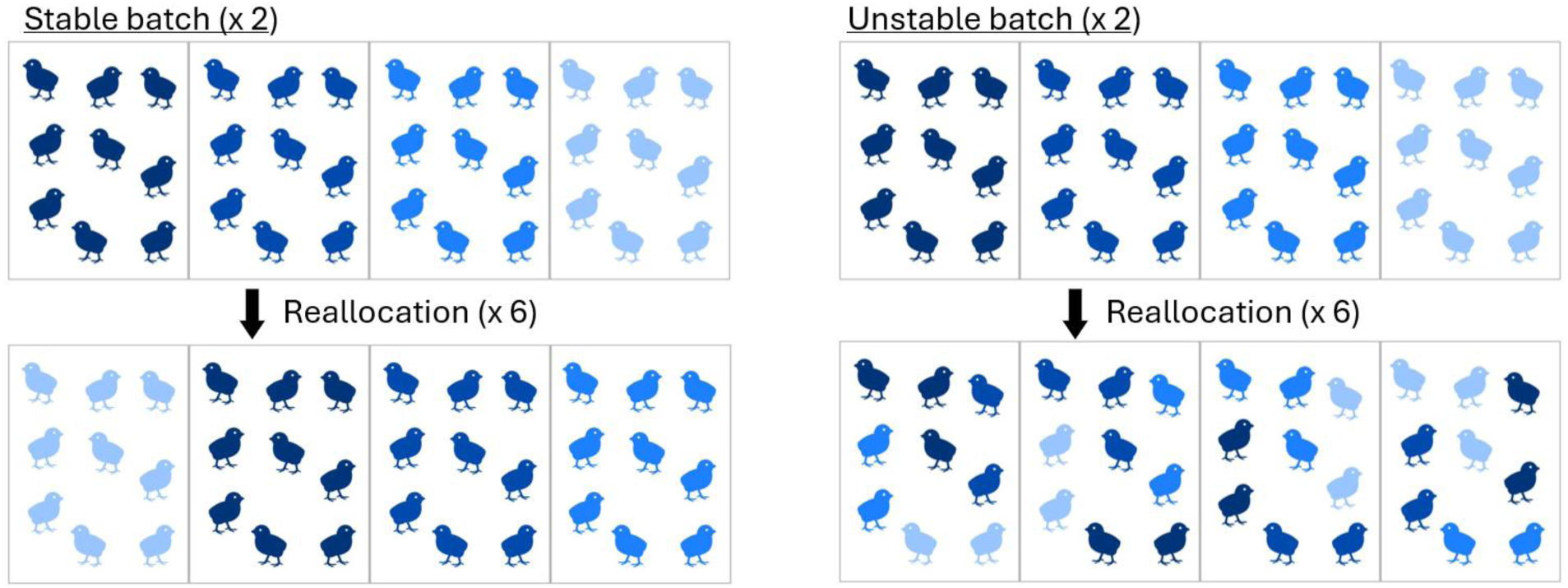
Schematic representation of the reallocation procedure for stable (left) and unstable (right) social treatments.

### Recording Aggression

Throughout the study, cameras positioned above the enclosures offered an overhead view of the birds. Recordings made from these cameras were used to assess the birds’ aggressive behaviour within their home enclosures. Our aim was to determine the impact of the social stability manipulation on the birds’ aggression. Therefore, we focused on the final day of the manipulation (day 28; Fig. 1) to capture the cumulative effect of the manipulation on the birds’ behaviour. We focussed on two ten-minute observation periods – the first immediately following the final reallocation, during which access to the feeder was partially restricted (see Supplementary Methods), and the second two hours later. The initial period was chosen as we expected it to be especially relevant to the birds’ aggressive behaviours, as we expected aggression to peak following the reallocation and restricted food access, and thus reveal key behavioural differences between the treatment groups. The additional observation two hours later allowed us to see if and how aggressive behaviour differed after more time since the reallocation had passed.

### Testing Response Inhibition

#### General Procedure and Test Arena

During the testing period (days 29 – 43; Fig. 1) the birds remained housed in the enclosures they had been moved to during the last reallocation on day 28.

Testing occurred during both morning (starting at 09:00) and afternoon (starting at 12:00) sessions to make it feasible to test all birds from a batch within the same day. Birds tested in the morning were food deprived the night before by removing the feeder at 17:00, and birds to be tested in the afternoon were food deprived in the morning by removing the feeder at 08:30. Additional analysis confirmed that the time of day that an individual was tested did not influence their performance (Table S1).

During testing, all individuals from an enclosure were caught and placed randomly into cat carriers in groups of three. The carriers were placed outside the testing room. One bird at a time was taken from a carrier, tested, and then placed back in the same carrier. Once all birds from an enclosure had been tested, they were all released back into their enclosure.

The test arena consisted of a wooden box (88 cm x 75.5 cm x 60 cm). It was lit with a single light bulb, attached next to a camera (BASCOM) mounted 80 cm above the centre of the arena. A tarpaulin ‘roof’ was pulled across the arena during testing to discourage and prevent birds from jumping out. A ‘start box’ (43.5 cm x 26 cm x 26.5 cm) was attached to the front of the testing arena, in which birds were placed before being let into the test arena. The start box had two ‘guillotine’ doors, one transparent and one opaque, and a separate light. Inside the start box was another square piece of wood attached to a handle, which could be used to gently push the birds into the test arena when required.

For each trial, the bird to be tested was first placed in the start box with both start box doors closed. First the opaque door was opened, allowing the bird to see but not enter the test box. After 10 s the transparent door was opened; the bird was allowed to remain in the start box for a further 15 s before being gently pushed into the test box. The end of the trial was indicated by the light in the test arena being switched off, and the light in the start box being switched on. The birds were allowed to return to the start box by themselves, or were gently guided back in when necessary. The start box doors would then be closed. The birds would be removed from the start box and placed back in their carrier at the end of the testing session.

### Habituation

First, the birds were habituated to the arena and general test procedures. On day 29, the birds were habituated in a group of three, formed at random from the nine birds currently sharing an enclosure. The stickers placed on the birds’ backs during the last reallocation remained, so that individuals could be differentiated in the video. On days 30, 31 and 32 the birds were habituated alone. The individual habituation procedure was repeated on day 36, which aimed to minimise any effect of the preceding thwarting task on birds’ food motivation on the subsequent cylinder task.

Group and individual habituation trials followed the same procedure. During each habituation trial, a large food dish (⌀ 25 cm) was placed in the test arena, containing the birds’ normal food (Wielink Kuikenmeel, Junai) as well as a handful of dried mealworms. Group habituation trials lasted for 8 mins, whereas individual habituation trials ended after 4 minutes.

### Thwarting

The thwarting task was administered to each bird twice, once on day 33 and once on day 43 (Fig. 1). This was done to evaluate whether birds changed their responses during this task after having experienced the cylinder task.

During the thwarting task, a barrier was placed centrally in the test arena (44.5cm from the front) and a food dish (same as used during habituation) was placed behind it (Fig. 3). The barrier consisted of a thin wood frame supporting a piece of chicken wire mesh. This filled the space such that the back half of the test arena was inaccessible to the chickens, with the barrier physically blocking access to the visible food. A thwarting trial lasted 4 minutes.

**Fig. 3.**
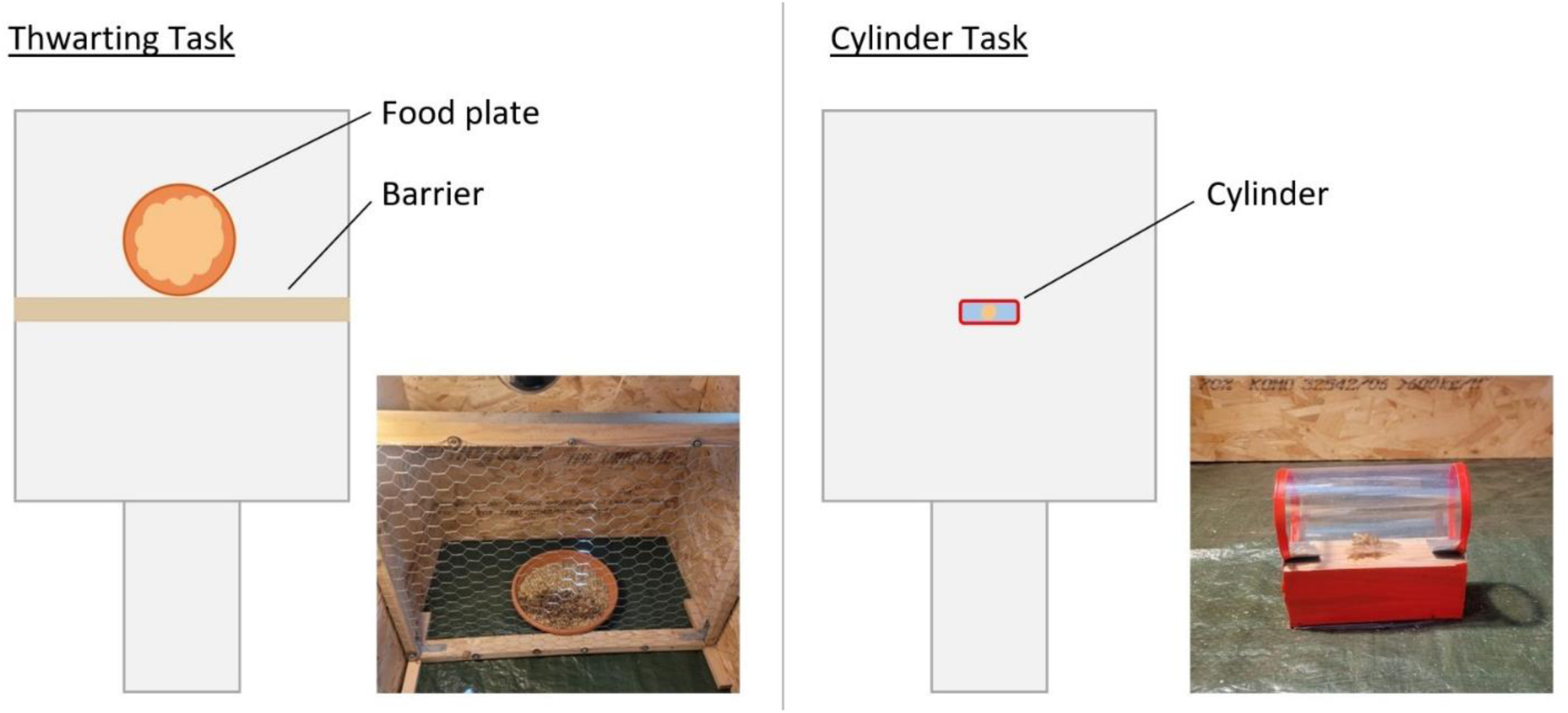
Schematics and photos of the apparatus used for the thwarting task (left) and the cylinder task (right).

### Cylinder

The cylinder task was administered across four days (days 37-40). Each bird received three trials on the first day, three trials on the second day, two trials on the third day and two trials on the fourth day, for a total of ten trials each.

A cylinder apparatus consisting of a transparent cylinder made of thin flexible plastic with open ends (⌀ 6 cm x 12.5 cm length) mounted on a wooden base (12.5 cm x 4.8 cm x 4.5 cm) was placed 41 cm from the entrance to the test arena so that the entrances were perpendicular to the start box, and secured to the floor with tape. The base was covered with red tape, and red tape was also placed around the entrances to the cylinder. A small pile of dried mealworms (∼ 1 tsp) was placed centrally in the cylinder (Fig. 3).

Each cylinder trial ended either when the bird ate from the cylinder, or after 2 mins. Between consecutive trials on the same day, the bird was guided back to the start box and remained there whilst the apparatus was re-set. After the last trial the bird returned to the start box, from which it was removed and placed back in its carrier.

### Video Coding and Experimental Variables

All videos were coded using the software BORIS (Behavioral Observation Research Interactive Software, version 9.2.1; Friard and Gamba 2016). All videos were coded by one experimenter (KW). A second person, naïve to the conditions, coded 20% of the videos (selected randomly) to assess inter-observer reliability. Cohen’s kappa coefficient for each of these videos was calculated within BORIS.

### Aggression

During coding, we focussed on aggressive behaviours that could be reliably identified from the top-down camera view (Table 1). Behaviour was continuously recorded for one enclosure at a time, using all-occurrence sampling. For each instance of aggression, we recorded the type and duration of the behaviour and the identities of the individuals involved. The average Cohen’s kappa value indicated a moderate to strong level of agreement between observers (k = 0.79, McHugh 2012).

**Table 1.**
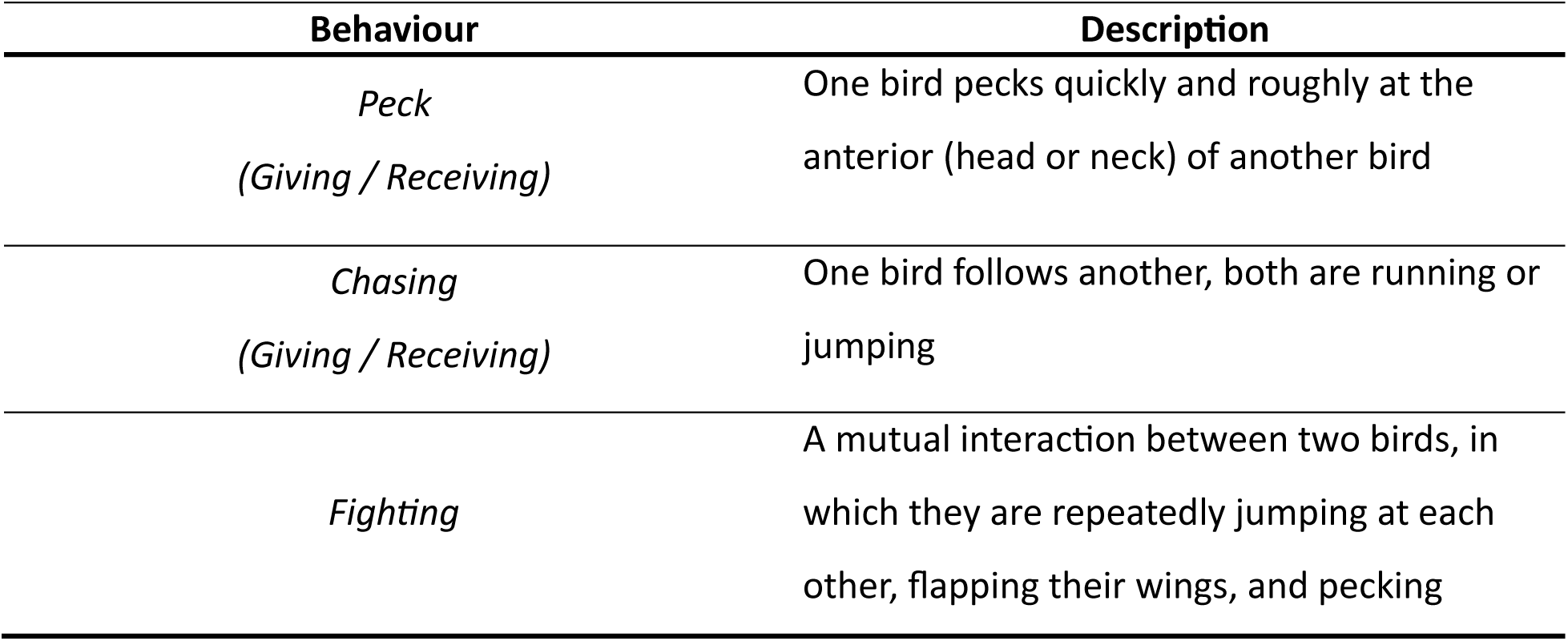
Ethogram of aggressive behaviours recorded during video coding (Arnould et al., 2024; Väisänen et al., 2005).

### Response Inhibition

In the thwarting task, we recorded the Time Spent Pecking, defined as the time the bird spent facing the barrier, actively and repeatedly pecking at it, for each bird in each trial. This duration paused if the bird faced away from the barrier or stopped pecking.

For each bird in each trial of the cylinder task, we recorded whether the bird was successful (i.e. the bird reached the food without first pecking the outside of the cylinder) or not. If the bird pecked within a trial, we also recorded the Time Spent Pecking, defined as the time the bird spends repeatedly actively pecking or pushing the closed sides of the cylinder in an attempt to reach the food. This duration paused if the bird faced away from the cylinder or stopped pecking. The average Cohen’s kappa value indicated a moderate to strong level of agreement between observers for the response inhibition tasks (Thwarting: k = 0.85, Cylinder: k = 0.79, McHugh 2012).

In both tasks, it was possible for a bird to not interact with the apparatus during a trial. If this was the case, they were not included when analysing the time spent pecking. We conducted an exploratory analysis to examine predictors of not interacting with the apparatus; details and results are provided in the Supplementary Information (Table S2, Table S3, Fig. S1, Fig. S2).

### Food motivation, body condition and sex

Individuals’ performance in response inhibition tasks has been shown to be affected by variation in motivation (van Horik et al. 2018; Willcox et al. 2024). As a proxy for the birds’ general motivation within the test box (which could be a combination of neophobia or neophilia, food motivation or motivation to explore; Dewulf et al. 2025), we recorded their latency to eat during their third individual habituation trial (day 32). This specific habituation trial was their last before testing began. This was done to ensure that the birds’ responses were less influenced by stress and not influenced by any prior testing.

Motivation to access the food rewards present in both tasks could also be influenced by the birds’ body condition (Shaw, 2017; Willcox et al. 2024). Therefore, the chickens’ body condition was calculated as weight/tarsus, from measurements taken on day 28 (i.e. end of the manipulation, prior to testing).

Finally, sex has been shown to influence response inhibition performance in other species (Junttila et al. 2021; Lucon-Xiccato 2022; van Horik et al. 2019; Willcox et al. 2024). The chicks were sexed by the hatchery, and this was later verified by visual inspection of the birds when they were 57 days old.

### Statistical analysis

All analyses were conducted in R (v. 4.5.1; R Core Team). Plots were created using the package *ggplot2* (Wickham et al. 2016).

### Aggressive Behaviour

We fitted a multivariate Bayesian linear mixed model using the package *brms* (v. 2.22.0; Bürkner 2021), including the three aggressive behaviours as dependent variables – number of pecks, time spent chasing and time spent fighting. Predictors were shared across sub-models and included the interaction between treatment group and observation time (either the first, immediately after reallocation, or the second, 2 hours later), with enclosure included as a random effect. Each response variable was modelled with its own appropriate likelihood (negative binomial distribution for pecks, hurdle gamma distribution for chasing and fighting).

To check model convergence and fit, we checked that all the Rhat values were close to 1 and that the ESS were above 1000, plotted density and trace of samples, and used graphical posterior predictive checks to assess model fit for each dependent variable. To examine the correlation structure between the three aggression measures at the enclosure level, we used the function *VarCorr* from *brms* to extract and interpret the estimated random effect variances and correlations. We used the *hypothesis* function in *brms* to compute post-hoc comparisons to test specific hypotheses about the interaction between group and observation effects. These included contrasts to test group differences within each observation time, as well as changes over time within each group.

### Response Inhibition

We fitted Bayesian general linear mixed models using *brms* on three dependent variables: time spent pecking in the thwarting task; success in the cylinder task; and time spent pecking in the cylinder task. All models included trial number, treatment group, sex, body condition and latency to eat (during habituation) as predictor variables, and individual ID as a random effect. Initial models also included the interaction between trial and group; when no evidence for an interaction was found, the interaction was removed. As the thwarting models provided no evidence for an effect of trial (see Results), we included the time spent pecking during the first trial of the thwarting task as a predictor variable in the cylinder models, to assess whether thwarting task performance predicted cylinder task performance. The predictors latency to eat and pecking during thwarting were log transformed to improve model stability. We used generic, weakly informative priors in all models. We used a lognormal family for both models looking at times spent pecking, and a Bernoulli family for the success model.

To check model convergence and fit, we checked that all the Rhat values were close to 1 and that the ESS were above 1000, plotted density and trace of samples and correlations between pairs, and used graphical posterior predictive checks to assess model fit. We used the *equivalence_test* function from the package *bayestestR* (v.0.15.2; Makowski et al. 2019) to assess the practical significance of the predictor variables, which evaluated how much of a predictor’s posterior distribution fell within the Region of Practical Equivalence (ROPE). Evidence for the null hypothesis is supported if 100% of the distribution fell within the ROPE, evidence to reject the null hypothesis is supported if 0% fell within the ROPE, and if only part of the distribution overlapped with the ROPE the evidence for the null hypothesis is undecided. We used post-hoc estimated marginal means tests using the package *emmeans* (v. 1.10.3; Lenth 2025) to investigate interactions. We also calculated bayes factors to provide further evidence regarding the effect of the predictor variables. Bayes factors were calculated by comparing the likelihood of the data under two models, one with the effect of interest included and one without, using the *bayes_factor* function from *brms*. A bayes factor above 1 indicates evidence in favour of the model with the effect, while a bayes factor below 1 indicates evidence for the null (Schmalz et al. 2023).

### Individual-level Aggression and Response Inhibition

To further investigate the potential link between response inhibition and aggression regulation, we conducted an additional exploratory analysis to see whether there was evidence that individual aggression profiles were associated with response inhibition performance, and whether these effects differed between treatment groups.

For this analysis, we sub-set the data to individuals that were involved in at least one instance of aggression (either as aggressor or recipient; n = 41 from stable groups, n = 53 from unstable groups). We then conducted a principal component analysis (PCA) using the packages *FactoMineR* (v. 2.12; Lê and Husson 2008) and *factoextra* (v. 1.0.; Kassambara and Mundt 2020) on the frequencies of the behaviours Giving Chase, Receiving Chase, Giving Peck, Receiving Reck and Fighting. All behavioural variables were standardised. The PCA was based on a correlation matrix. Only components with Eigenvalues higher than 1 were retained, and for each component we only considered loadings whose contribution was higher than what was expected if all loadings were of equal weight.

To examine whether individual-level aggression was associated with response inhibition, we conducted Spearman’s rank correlations between aggression scores (PC1 and PC2) and the response inhibition measures (time spent pecking in the first trial of the thwarting task, total number of successful trials in the cylinder task, and mean time spent pecking in the cylinder task).

## Results

### Aggressive Behaviour

The average (±SE) number of pecks, time spent chasing and time spent fighting across treatment groups and observations are visualised in Figure 4. We found evidence for an interaction between observation and treatment group for chasing and fighting, but not pecking (Table 2). Post-hoc comparisons provided evidence that there was more aggression observed in unstable groups than in stable groups during the first observation (right after the reallocation), but that the groups did not differ during the second observation (Table 3; Fig. 4). We found evidence that all three aggressive behaviours decreased from the first to the second observation in unstable groups, but that only pecking decreased in the stable groups (Table 4). We found evidence for high variation in aggression between enclosures (Table S4), and did not find evidence for correlations between different aggression behaviours at the enclosure level (Table S5).

**Fig. 4.**
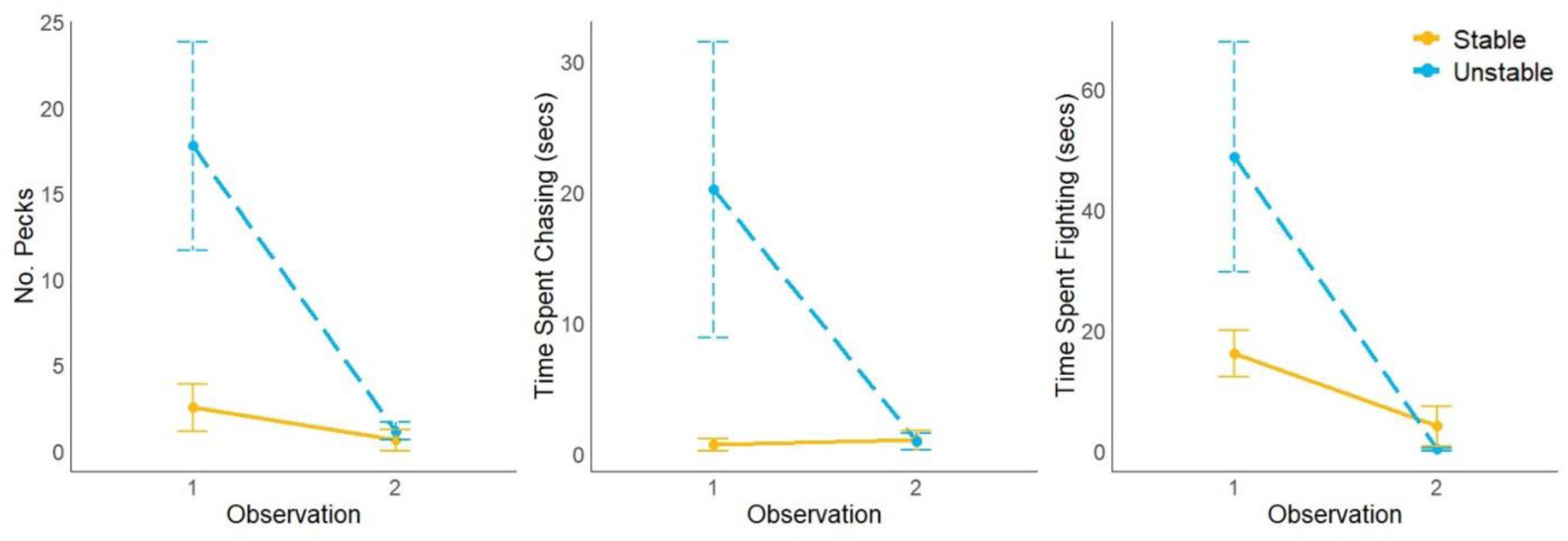
The average (± SE) number of pecks (left), time spent chasing (middle) and time spent fighting (right) observed in enclosures, in stable and unstable groups across two observations: 1) immediately following the final reallocation and 2) two hours later.

**Table 2.** Estimates and credible intervals (CI) from the multivariate model investigating the influence of the interaction between group and observation time on aggression. Results indicating evidence for an effect of the interaction (95% credible interval not crossing 0) are highlighted in bold.

| Parameter | Dependent Variable | Estimate | 95% CI low | 95% CI up |
| --- | --- | --- | --- | --- |
| Group * Observation | Pecks | -0.77 | -2.59 | 1.12 |
|  | **Chasing** | **-3.35** | **-5.70** | **-0.95** |
|  | **Fighting** | **-2.80** | **-4.37** | **-1.14** |

**Table 3.**
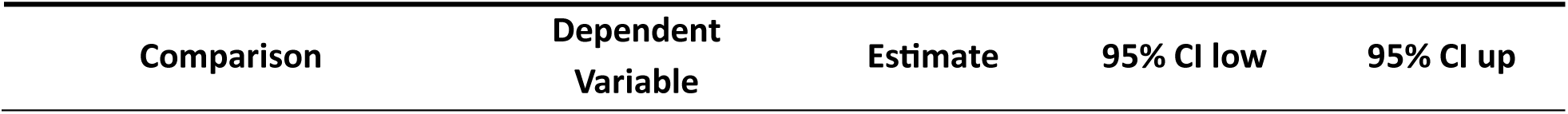
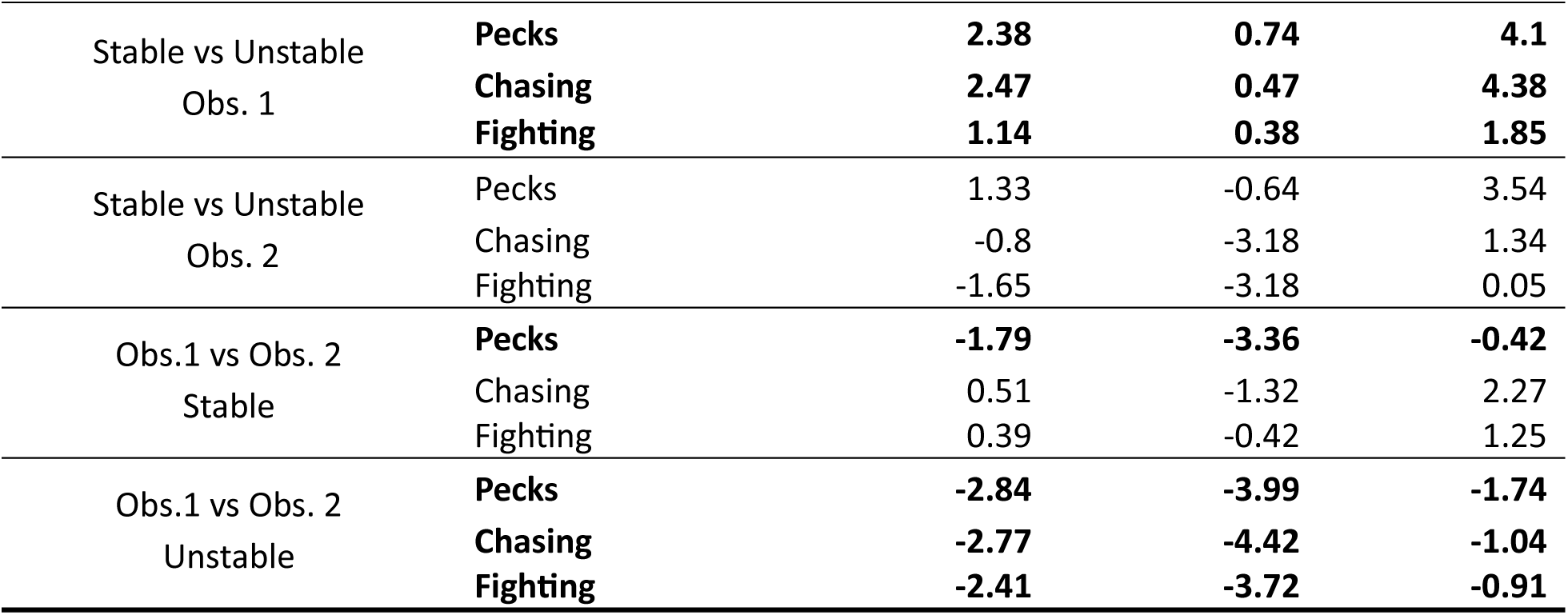
Estimates and credible intervals (CI) from the post-hoc tests investigating the influence of the interaction between group and observation (Obs.) on aggression. Results indicating evidence for a difference between the compared parameters (95% credible interval not crossing 0) are highlighted in bold.

**Table 4.**
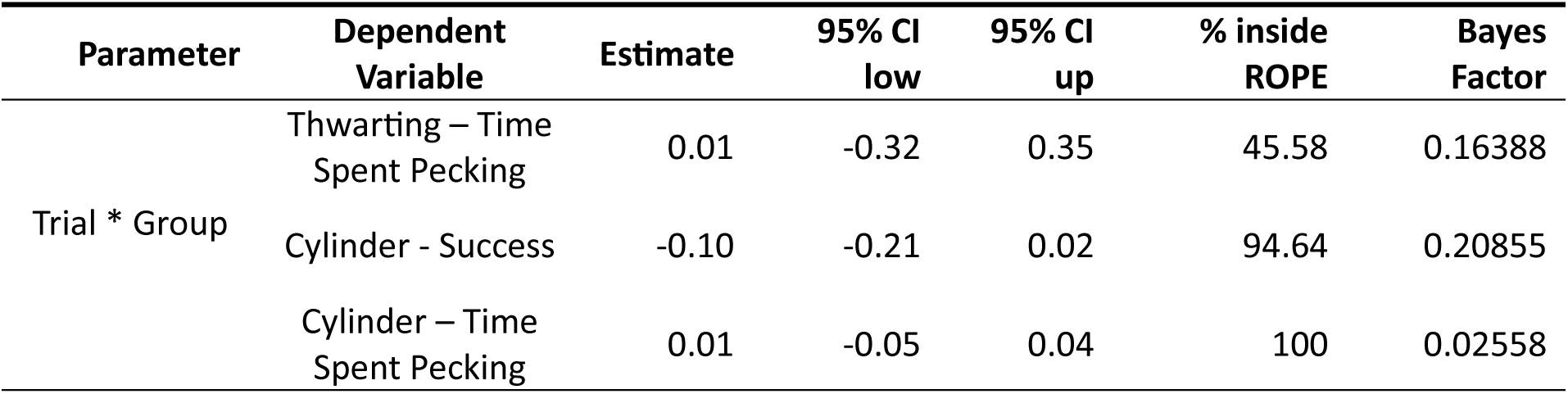
Estimates and test statistics from the post-hoc EMMs tests investigating the influence of the interaction between trial and group on performance in the Thwarting and Cylinder Tasks. CI – Credible Interval, ROPE – Region Of Practical Equivalence.

| Parameter | Dependent Variable | Estimate | 95% CI low | 95% CI up | % inside ROPE | Bayes Factor |
| --- | --- | --- | --- | --- | --- | --- |
| Trial * Group | Thwarting – Time Spent Pecking | 0.01 | -0.32 | 0.35 | 45.58 | 0.16388 |
|  | Cylinder - Success | -0.10 | -0.21 | 0.02 | 94.64 | 0.20855 |
|  | Cylinder – Time Spent Pecking | 0.01 | -0.05 | 0.04 | 100 | 0.02558 |

### Response Inhibition

*Thwarting*.

We found no evidence for an influence of an interaction between trial and treatment group for time spent pecking (Table 4). After removing the interaction, the simpler model produced no evidence of an effect of trial, group, sex, body condition or latency to eat on time spent pecking (Table 5, Fig. 5).

**Fig. 5.**
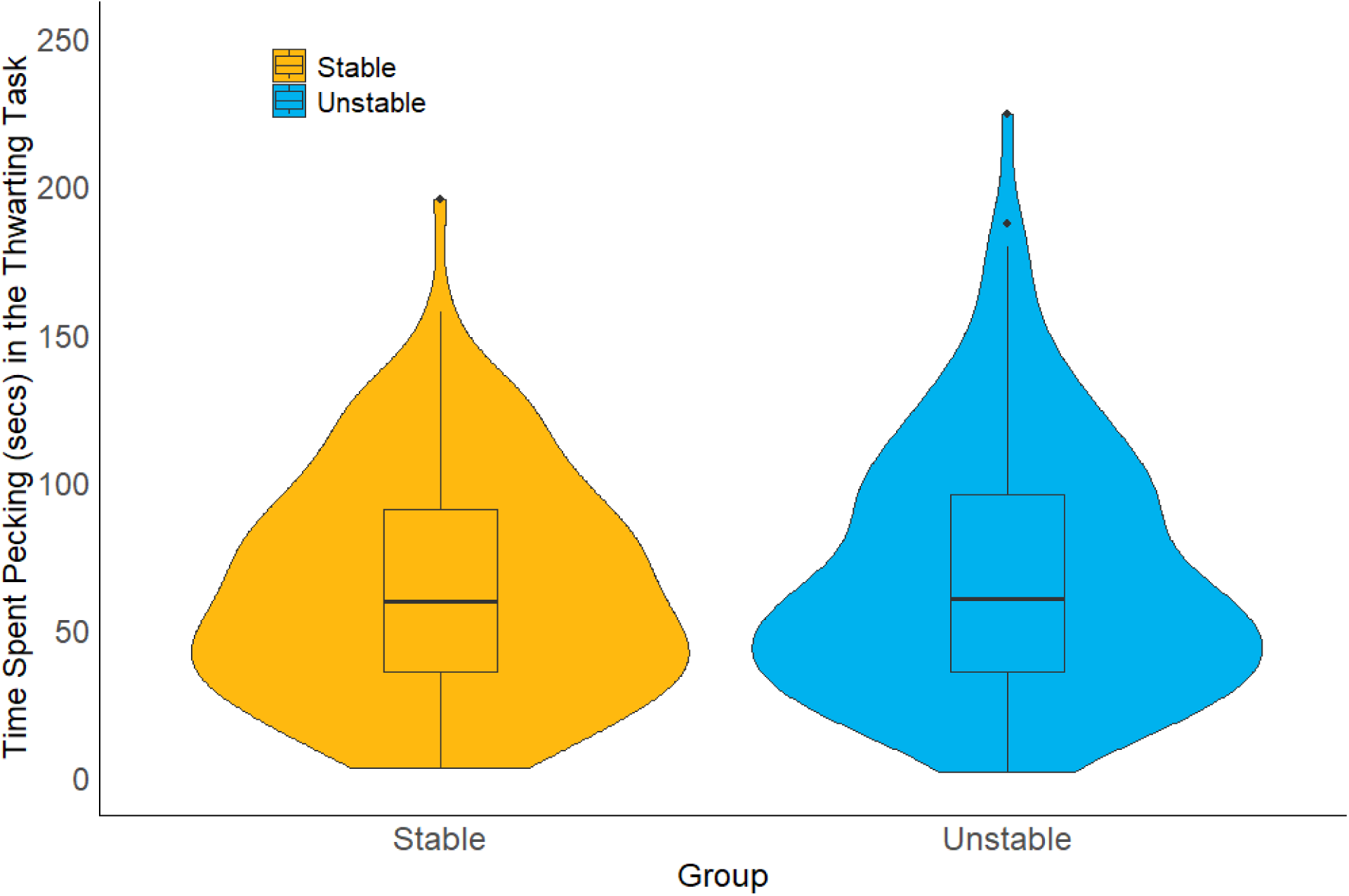
Time spent pecking in the thwarting task by chickens raised in stable and unstable groups that pecked. Includes data collated across two trials.

**Table 5.** Estimates and test statistics from the models assessing performance in the Thwarting and Cylinder Tasks. CI – Credible Interval, ROPE – Region Of Practical Equivalence. Results indicating evidence for an effect of the parameter (95% credible interval not crossing 0, 0% of posterior distribution falling within the ROPE, Bayes Factor > 1) are highlighted in bold.

| *Thwarting Task – Time Spent Pecking* |  |  |  |  |  |
| --- | --- | --- | --- | --- | --- |
| Parameter | Estimate | 95% CI low | 95% CI up | % inside ROPE | Bayes Factor |
| Trial | -0.16 | -0.33 | 0.02 | 2.19 | 0.43790 |
| Group | 0.11 | -0.08 | 0.30 | 4.69 | 0.19524 |
| Sex | 0.13 | -0.06 | 0.32 | 3.42 | 0.24137 |
| Body Condition | -0.05 | -0.13 | 0.02 | 8.25 | 0.11677 |
| Latency to Eat | 0.04 | -0.06 | 0.14 | 11.68 | 0.07413 |
| *Cylinder Task - Success* |  |  |  |  |  |
| Parameter | Estimate | 95% CI low | 95% CI up | % inside ROPE | Bayes Factor |
| **Trial** | **0.31** | **0.25** | **0.38** | **0** | **> 1000** |
| Group | -0.29 | -0.71 | 0.12 | 28.52 | 0.55279 |
| Sex | -0.06 | -0.47 | 0.35 | 63.39 | 0.22009 |
| Body Condition | -0.06 | -0.22 | 0.09 | 95.46 | 0.10601 |
| Latency to Eat | 0.05 | -0.17 | 0.26 | 91.29 | 0.11751 |
| Pecking in Thwarting Task | 0.07 | -0.09 | 0.23 | 94.98 | 0.07342 |
| *Cylinder Task – Time Spent Pecking* |  |  |  |  |  |
| Parameter | Estimate | 95% CI low | 95% CI up | % inside ROPE | Bayes Factor |
| **Trial** | **-0.10** | **-0.12** | **-0.08** | **0** | **> 1000** |
| Group | -0.10 | -0.24 | 0.04 | 4.35 | 0.18598 |
| Sex | -0.09 | -0.23 | 0.05 | 5.10 | 0.16835 |
| Body Condition | 0.02 | -0.03 | 0.07 | 24.75 | 0.03218 |
| Latency Eat | -0.01 | -0.08 | 0.06 | 21.82 | 0.03860 |
| Pecking in Thwarting Task | -0.06 | -0.13 | < -0.01 | 2.70 | 0.20048 |

### Cylinder

We found no evidence that the change in success across trials differed between treatment groups (Table 4). After removing the interaction, the simpler model provided very strong evidence that success increased across trials (Table 5, Fig. 6), but no evidence for an effect of group, sex, body condition, latency to eat or time spent pecking in the thwarting task.

**Fig. 6.**
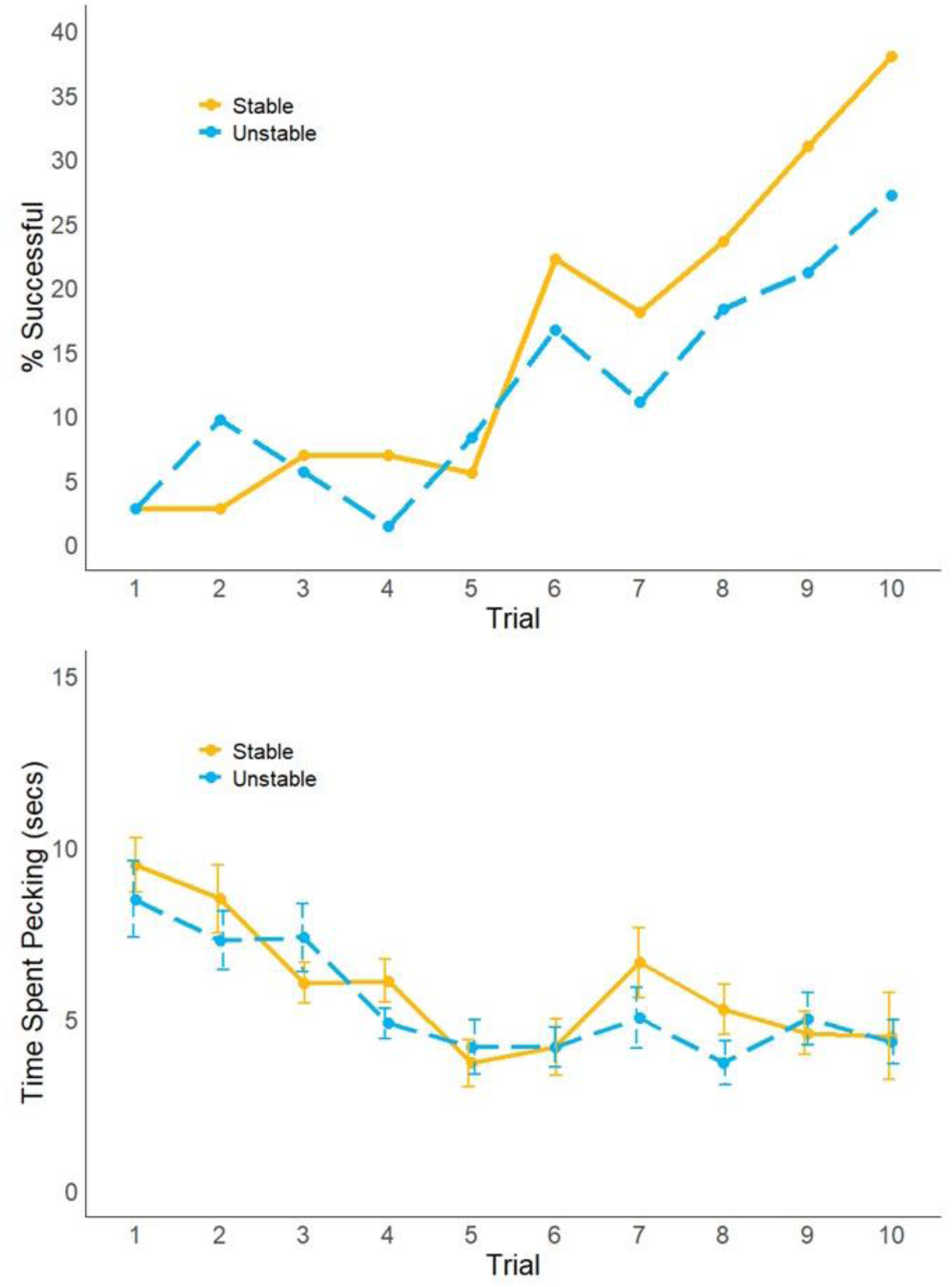
Performance in the cylinder task: above: the percentage of chickens from stable and unstable groups that were successful in each trial of the cylinder task; below: the time spent pecking (mean ± SE) by chickens raised in stable and unstable groups that pecked, for each trial of the cylinder task.

We found no evidence that the change in time spent pecking across trials differed between treatment groups, instead we found strong evidence for the null hypothesis (Table 4). After removing the interaction, the simpler model provided very strong evidence that time spent pecking decreased across trials (Table 5, Fig. 6), and no evidence for an effect of group, sex, body condition, latency to eat or time spent pecking in the thwarting task.

### Individual-level Aggression and Response Inhibition

A total of 94 birds were involved in at least one instance of aggression, either as aggressor or recipient; 41 from stable groups and 53 from unstable groups. In the PCA, behaviours related to being aggressive primarily loaded on PC1, while behaviours related to receiving aggression primarily loaded on principal component 2 PC2 (Table S6). We found no correlation between the aggression scores (PC1 and PC2) and the response inhibition measures (time spent pecking in the first trial of the thwarting task, total number of successful trials in the cylinder task, and mean time spent pecking in the cylinder task, Fig. S3).

## Discussion

We predicted that early-life social instability would alter demands for aggression regulation, influencing the development of response inhibition in chickens. We observed more aggression in unstable groups immediately following the final reallocation, but aggression decreased to the same lower level as in the stable groups within two hours. In contrast to our predictions, we found that performance in two response inhibition tasks did not differ between birds raised in stable groups compared to unstable groups. Further analysis revealed that individual-level involvement in aggressive interactions was not associated with response inhibition performance. Together, these results suggest that early-life social instability influences social behaviour, but not response inhibition, in chickens.

Our observations of the birds in their enclosures revealed that more aggression occurred in unstable groups than in stable groups, demonstrating that the manipulation influenced the birds’ social behaviour. This is consistent with previous observations that chickens interact more with unfamiliar than with familiar individuals, with both aggressive and affiliative, explorative social interactions (Riedstra and Groothuis 2002; Parada et al. 2021; Arnould et al. 2024). Initial aggression between unfamiliar individuals may function to establish a dominance hierarchy, which reduces aggression once stable (Pagel and Dawkins 1997, Väisänen et al. 2004; Desire et al. 2015). The rapid decline in aggression in unstable groups could, therefore, show an efficient establishment of new dominance hierarchies. Alternatively, the reduction could reflect social tolerance. Broiler chickens have been found to be more socially tolerant than layer strains (Mench 1988), and in very large groups, they adopt a strategy of reduced aggression (instead of hierarchy formation) to cope with repeatedly encountering unfamiliar individuals (Estevez 1997). The tendency for such social tolerance in broilers may have facilitated the quick decline in aggression following mixing. The pattern of aggression observed in unstable groups indicates a rapid reduction in aggressive behaviour, thereby avoiding the costs of prolonged conflict.

Previous studies have shown response inhibition to be influenced by social complexity (Amici et al. 2008; Amici et al. 2018; Joly et al. 2017; Loyant et al. 2023; Ashton et al. 2018; Johnson-Ulrich and Holekamp 2020; Vernouillet et al. 2025; Willcox et al. 2024), including an influence of social stability specifically (Lucon-Xiccato et al. 2022; Amici et al. 2008). However, despite using a controlled social manipulation with a large sample size, and two different tasks assessing different aspects of response inhibition, we found no evidence that early-life social stability influenced performance in either task. In the cylinder task, the birds learned to inhibit their actions, as evidenced by an increase in successful trials and a decrease in the time spent pecking (Vernouillet et al. 2016, Willcox et al. 2024), but learning trajectories did not differ between the treatment groups. Our findings suggest that early-life social stability does not shape response inhibition, nor the learning of it, in chickens, questioning the generalisability of previously reported links between social complexity and response inhibition.

Together, these results suggest that the group differences in aggression did not translate into corresponding group-level differences in response inhibition. Further analysis also revealed that individuals’ aggression profiles were not associated with their inhibition performance, in contrast to previous findings (Vernouillet et al. 2025; Overduin-de Vries et al. 2023, Gobbo and Semrov et al. 2022; Rudebeck et al. 2007; Cervantes and Deville, 2007). This could suggest that although social instability influenced aggression, differences in the *demands for aggression regulation* between the two treatment groups were not sufficient to drive variation in response inhibition. This may reflect the sociality of our study species; broiler chickens are relatively socially tolerant (Mench 1988; Estevez 1997) and form new dominance hierarchies rapidly after mixing, as supported by our aggression results. Consequently, the unstable environment may not have imposed strong or prolonged demands requiring either enhanced or reduced inhibition. In less tolerant species, such as the guppies tested by Lucon-Xiccato et al. (2022), continued aggression may remain advantageous for resource access and potentially favour reduced response inhibition. Conversely, in species where aggression could carry long-term social costs, such as primates with fission-fusion societies (Amici et al. 2008; Amici et al. 2018; Aureli et al. 2008), the demands for restraint may be greater in unstable environments, promoting stronger inhibition.

It is also possible that relevant aspects of social behaviour went unmeasured in our study. Our behavioural observations were limited to overt aggression visible in top-down video recordings, but chickens also engage in more subtle dominance-related behaviours such as threats (Arnould et al. 2024; Väisänen et al. 2005), and affiliative social behaviours (Riedstra and Groothuis 2002; Parada et al. 2021), that could also be relevant for social demands on response inhibition. A more comprehensive approach to characterising social behaviour, potentially using social network analysis, would help capture the full complexity of social environments and their cognitive implications (Wascher et al. 2018; Speechley et al. 2024a).

Another potential explanation for our finding that early-life social stability did not influence response inhibition is our choice of tasks. We used two tasks that were similar in that they both included the same “go” stimulus (visible food) and a similar “stop” stimulus (impenetrable barrier) (Troisi et al. 2025), yet differed in their physical characteristics (including the size of the go stimulus), and the accessibility of the reward. We found no relationship between individual’s performance on the thwarting task and the cylinder task, similarly to previous studies showing a lack of repeatability in individual’s performance across response inhibition tasks (Bray et al. 2013; Marshall-Pescini et al. 2015; Brucks et al. 2017; Vernouillet et al. 2018; van Horik et al. 2018; Szabo et al. 2020; Loyant et al. 2022; Vernouillet et al. 2025; Troisi et al. 2025). This suggests that these tasks did measure different aspects of response inhibition. Measures in different tasks could be differentially influenced by early-life social environments, potentially accounting for conflicting findings in previous studies. Despite this, we found no influence of social stability on measures in either task. Dunbar and Shultz (2025) argue that cylinder-type tasks are more closely tied to physical or foraging-related inhibition, while tasks requiring “strategic self-control” (e.g. A-not-B or delayed gratification paradigms) show stronger links to social demands; perhaps such tasks would better reflect the inhibition potentially involved in regulating aggression. However, this cannot account for the findings of previous studies that reported an effect of the social environment on response inhibition using cylinder or thwarting-type tasks (Ashton et al. 2018; Johnson-Ulrich and Holekamp 2020, Vernouillet et al. 2025, Lucon-Xiccato et al. 2022).

Crucially, both the thwarting and cylinder tasks lacked social components. As we did find an effect of early-life social stability on aggression, it is possible that the manipulation had effects that are specific to social contexts and thus not captured by these tasks. This raises the broader question of whether social environments exert generalised effects on cognition, or whether their impact is domain-specific. Early-life social experiences have been shown to affect performance in non-social tasks in several studies (Lambert and Guillette 2021; Speechley et al. 2024b; Szabo and Ringler 2025), including those testing response inhibition (Ashton et al. 2018; Johnson-Ulrich and Holekamp 2020, Vernouillet et al. 2025, Lucon-Xiccato et al. 2022). However, other findings indicate domain-specific effects, in which social environments influence specifically social, but not non-social, behaviour and cognition (Ferreira et al. 2025; MacLean et al. 2013; reviewed in Taborsky and Oliverira 2012). Our findings are consistent with the concept of domain-specificity as we found an influence of the social environment on social behaviour, but not cognition in a non-social context. Future studies would benefit from testing cognition and behaviour in a variety of social and non-social contexts, and in species with varying social systems, to gain a more comprehensive understanding of the influence of early-life social environments.

Unlike some previous studies, we found no evidence that variation in individual factors influenced the birds’ performance. Despite motivation to interact with an apparatus (driven by food motivation and/or reduced neophobia) having been found to influence individuals’ response inhibition performance in other bird species (Shaw 2017; Willcox et al. 2024; Vernouillet et al. 2025), here we found no influence of the chickens’ body condition or a proxy of their general motivation (latency to eat during habituation) on any of the response inhibition measures. It is possible that our controlled raising and habituation procedures reduced individual variation in these traits. In other species, sex has also been shown to influence response inhibition, with females generally showing better inhibition than males (Willcox et al. 2024, van Horik et al. 2019; Junttila et al. 2021; Lucon-Xiccato et al. 2022), a trend potentially driven by males being generally more ‘persistent’ due to the requirements of their mating behaviour (Vinogradov et al. 2022). We did not replicate this finding in chickens, despite male chickens being more persistent (Rogers 1974), and the sexes differing in other measures of their behaviour and cognition (e.g. Nätt et al. 2014; Vallortigara 1996; Santolin 2020), including in this same cohort of birds (albeit when they were older - Moldovan et al., in prep). This may be because the chickens were still young (and not sexually mature) during this study. Supplementary analysis did, however, reveal that female chickens were more likely to not interact with the apparatus in the thwarting task than males, potentially indicating some sex-specific differences in motivation and persistence in the chickens at this age.

## Conclusion

In summary, we found experimental evidence that early-life social instability temporarily increased aggression in groups of chickens, but that this did not influence their response inhibition. These findings question the role of response inhibition in aggression regulation, and challenge the idea that response inhibition is shaped by social experiences, at least for this species. Instead, they add to a growing literature suggesting that the effects of social complexity on behaviour and cognition are highly context- and species-dependent. Different measures and manipulations of social complexity are likely to place different demands on cognition depending on species-specific sociality, and tasks used to assess cognition may only reflect responses in the tested context. Future work should, therefore, combine multiple measures of cognition and behaviour, both social and non-social, and consider the ecology of the tested species, to gain a more comprehensive understanding of how early-life environments shape behavioural and cognitive development.

## Declarations

### Ethics Approval

The experiment was performed in accordance with the Association for the Study of Animal Behaviour ethical guidelines under permission of the ethical committee of animal experimentation (VIB Site Ghent, Universiteit Gent): EC2023-064.

### Data Availability

The raw data used in this study, and the code used to process and analyse them, are available from the Open Science Framework repository: https://osf.io/u9at8/overview?view_only=b20ebd09f4cf46c89472ba63d3d61592

## Acknowledgements

We thank Nathan Audenaert, Simon Braem, Pauline Blerot and Lumi Vanhulle for assisting with collecting the data for this study. We also thank Nathan Audenaert and Emilia De Graeve for assisting with validation, Vitor Ferreira for his advice during manuscript preparation, and the staff of the Wildlife Rescue Centre Ostend for general project support.

## Author Contributions

Based on the CRediT system, the contributions of the authors are as follows:

‘Conceptualization’ – KW, AV, LL, FV; ‘Methodology’ - KW, AV, LL, FV; ‘Validation’ – KW, BS; ‘Formal Analysis’ – KW; ‘Investigation’ - KW; ‘Resources’ – FV, AM; ‘Data Curation’ – KW; ‘Writing – Original Draft’ – KW; ‘Writing – Review & Editing’ – KW, AV, BS, AM, LL, FV; ‘Visualization’ – KW; ‘Supervision’ – AV, LL, FV; ‘Project Administration’ – KW, FV; ‘Funding Acquisition’ – AV, AM, LL, FV.

