## Supplementary for "Early-life social instability affects aggression, but not response inhibition, in chickens"

### Supplementary Information for 'Early-life social instability affects aggression, but not response inhibition, in chickens'

Kathryn Willcox, Alizée Vernouillet, Birgit Szabo, An Martel, Luc Lens and Frederick Verbruggen

Corresponding author: Kathryn Willcox (Centre for Research on Ecology, Cognition and Behaviour of Birds, Ghent University; Department of Experimental Psychology, Ghent University; Department of Biology, Ghent University),

#### Supplementary Methods

##### *Reallocation Procedure*

The reallocation procedure went as follows: the birds were first caught and placed in labelled cat-carriers in groups of two or three individuals (three carriers with two birds and one carrier with three birds, from each enclosure of nine). The carriers were then placed outside the enclosures. During this time, any other necessary activity would take place, such as replacing the birds' stickers, replacing the birds leg rings, and taking measurements (see above). If nothing was necessary, each bird would still be handled, such that this experience remained consistent throughout. Additionally, all the materials from the enclosures were cleaned during this time, and the substrate was replaced when necessary.

Once these steps were complete, the birds were placed back into their new enclosures. To do this, first all the materials were placed back into the enclosures, except for the feeders. Then four cat-carriers containing nine birds were placed into each enclosure, following the pre-determined reallocation plan. We opened the carriers, releasing the birds into the enclosure as simultaneously as possible. Then we replaced the feeders and shut the doors to the enclosures. At this time, all but two of the openings of each feeder were covered with grey tape, such that the food could only be accessed via the two open holes. After 15 minutes, we opened the doors, removed the tape from the feeders, and placed them back.

#### Supplementary Analysis & Results

*Did the time of day that birds were tested influence their response inhibition performance?*

To make it feasible to test all birds from one batch within the same day, testing for the cylinder task occurred in both morning (starting at 9:00) and afternoon (starting at 12:00) sessions. Birds tested in the morning were food deprived the night before by removing the feeder at 17:00, and birds to be tested in the afternoon were food deprived in the morning by removing the feeder at 8:30.

To check whether testing (and thus food deprivation) time impacted birds' performance in the cylinder task, we fitted Bayesian general linear mixed models using *brms* on success and time spent pecking in the cylinder task. Models included testing time (in the morning or in the afternoon) as a predictor variable, and individual ID as a random effect. We used generic, weakly informative priors in all models. We used a lognormal family for the time spent pecking model, and a Bernoulli family for the success model. Model checking and result extraction followed the same procedures as for the main response inhibition analyses.

We found no evidence that the time of day when birds were tested influenced their performance in the cylinder task (Table S1).

**Table S1.** Estimates and test statistics from the models investigating the influence of testing time on performance in the Cylinder Task. CI – Credible Interval, ROPE – Region Of Practical Equivalence.

| Parameter | Dependent Variable | Estimate | 95% CI low | 95% CI up | % inside ROPE | Bayes Factor |
|---|---|---|---|---|---|---|
| Testing Time | Cylinder - Success | -0.11 | -0.47 | 0.25 | 63.52 | 0.36965 |
| | Cylinder – Time Spent Pecking | -0.03 | -0.16 | 0.10 | 11.75 | 0.07399 |

*Birds did not always interact with the tasks – what predicted whether they did not interact?*

On each trial of both the cylinder and thwarting tasks, some birds did not interact with the apparatus. For the thwarting task, this occurred if the bird did not peck the mesh barrier at all during the trial. For the cylinder task, this occurred if the bird both did not peck the cylinder and also did not retrieve the reward. Especially with experience of the task(s), not interacting with the apparatus could also indicate response inhibition (i.e., inhibiting the ineffective behaviour of pecking the apparatus – Willcox, et al., 2024). Therefore, we assessed whether not interacting was influenced by treatment group, trial number, or any of the individual-level predictor variables.

We fitted Bayesian general linear mixed models using *brms* on two dependent variables: not interacting in the thwarting task, and not interacting in the cylinder task. All models included trial number, treatment group, sex, body size and latency to eat (during habituation) as predictor variables, and individual ID as a random effect. Initial models also included the interaction between trial and group; when no evidence for an interaction was found, the interaction was removed. The predictor latency to eat was log transformed to improve model stability. We used generic, weakly informative priors, and a Bernoulli family in all models.

We found no evidence for an interaction between trial and treatment group for not interacting in either the thwarting or cylinder task (Table S2). After removing the interactions, simpler models produced some evidence that not interacting in the thwarting task was influenced by sex, and that strong evidence that not interacting in the cylinder task was influenced by trial number, but no evidence that any of the other predictor variables had an effect (Table S3).

In the thwarting task, more female chickens did not interact with the apparatus than males (Fig S1). In the cylinder task, the number of birds not interacting with the apparatus decreased across trials (Fig. S2).

**Table S2.** Estimates and test statistics from the post-hoc EMMs tests investigating the influence of the interaction between trial and group on not interacting in the Thwarting and Cylinder Tasks. CI – Credible Interval, ROPE – Region Of Practical Equivalence.

| Parameter | Dependent Variable | Estimate | 95% CI low | 95% CI up | % inside ROPE | Bayes Factor |
|---|---|---|---|---|---|---|
| Trial * Group | Not Interacting in the Thwarting Task | 0.35 | -1.33 | 0.60 | 12.86 | 0.61866 |
| | Not Interacting in the Cylinder Task | 0.09 | -0.22 | 0.04 | 55.53 | 0.16140 |

**Table S3.** Estimates and test statistics from the models assessing not interacting in the Thwarting and Cylinder Tasks. CI – Credible Interval, ROPE – Region Of Practical Equivalence. Results indicating evidence for an effect of the parameter (95% credible interval not crossing 0, 0% of posterior distribution falling within the ROPE, Bayes Factor > 1) are highlighted in bold.

| *Not Interacting in the Thwarting Task* | | | | | |
|---|---|---|---|---|---|
| **Parameter** | **Estimate** | **95% CI low** | **95% CI up** | **% inside ROPE** | **Bayes Factor** |
| Trial | -0.32 | -1.12 | 0.46 | 27.86 | 0.56139 |
| Group | 0.42 | -0.39 | 1.23 | 22.25 | 0.67473 |
| Sex | -1.06 | **-1.95** | **-0.23** | **0** | **8.56205** |
| Body Size | -0.19 | -0.53 | 0.15 | 46.68 | 0.32953 |
| Latency to Eat | 0.28 | -0.12 | 0.67 | 30.01 | 0.54284 |
| *Not Interacting in the Cylinder Task* | | | | | |
| **Parameter** | **Estimate** | **95% CI low** | **95% CI up** | **% inside ROPE** | **Bayes Factor** |
| Trial | -0.23 | **-0.29** | **-0.16** | 6.63 | **> 1000** |
| Group | 0.60 | -0.30 | 1.52 | 14.12 | 0.19059 |
| Sex | -0.26 | -1.14 | 0.62 | 28.16 | 0.08820 |
| Body Size | 0.27 | -0.12 | 0.65 | 31.04 | 0.10072 |
| Latency to Eat | 0.54 | 0.06 | 1.03 | 5.21 | 0.52857 |

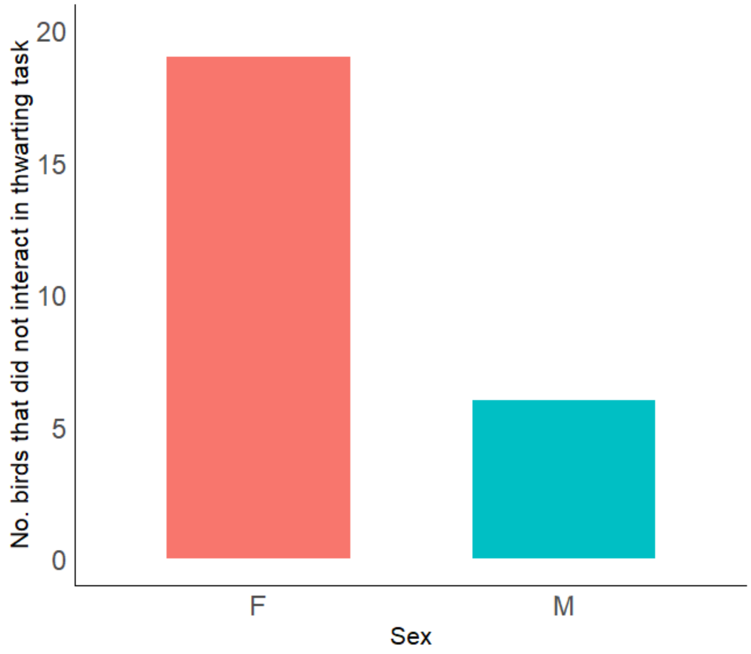


**Fig. S1** The number of female (F) and male (M) birds that did not interact with the apparatus in the thwarting task. Includes data collated across two trials

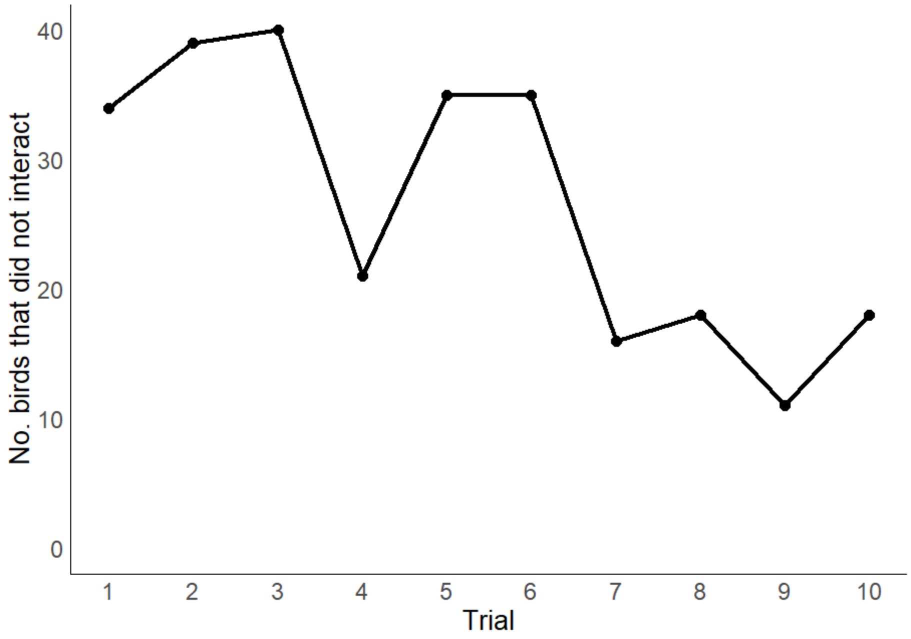


**Fig. S2** The number of birds that did not interact in each trial of the cylinder task

*Aggression*

**Table S4.** Posterior estimates of between-enclosure variation (SD of random intercepts) for each aggressive behaviour.

| Dependent Variable | Standard Deviation Estimate | 95% CI low | 95% CI up |
|---|---|---|---|
| Pecks | 1.26 | 0.53 | 2.27 |
| Chasing | 0.84 | 0.04 | 2.14 |
| Fighting | 0.50 | 0.07 | 0.98 |

**Table S5.** Posterior estimates and 95% credible intervals for correlations between random intercepts of aggressive behaviours at the enclosure level.

| Pair | Correlation Estimate | 95% CI low | 95% CI up |
|---|---|---|---|
| Pecks & Chasing | -0.06 | -0.84 | 0.76 |
| Pecks & Fighting | 0.06 | -0.67 | 0.71 |
| Chasing & Fighting | -0.02 | -0.86 | 0.85 |

*Individual-level Aggression*

**Table S6**. PCA loadings for the aggressive behaviours. Loadings considered important (i.e. whose contribution was higher than what was expected if all loadings were of equal weight) are indicated in bold.

| | **Dimensions** | | | | |
|---|---|---|---|---|---|
| | 1 | 2 | 3 | 4 | 5 |
| Eigen values | **2.26** | **1.13** | 0.73 | 0.69 | 0.19 |
| Variance explained (%) | 45.38 | 22.5 | 14.58 | 13.78 | 3.76 |
| Cumulative variance explained (%) | 45.38 | 67.88 | 82.46 | 96.24 | 100 |
| Chasing (Giving) | **0.61** | **-0.28** | 0.56 | 0.47 | -0.01 |
| Chasing (Receiving) | 0.129 | **0.88** | -0.19 | 0.44 | 0.02 |
| Pecks (Giving) | **0.89** | -0.16 | -0.03 | -0.03 | 0.31 |
| Pecks (Receiving) | **0.54** | **0.49** | 0.43 | -0.52 | 0,01 |
| Fighting | **0.89** | -0.08 | -0.32 | -0.04 | -0.31 |

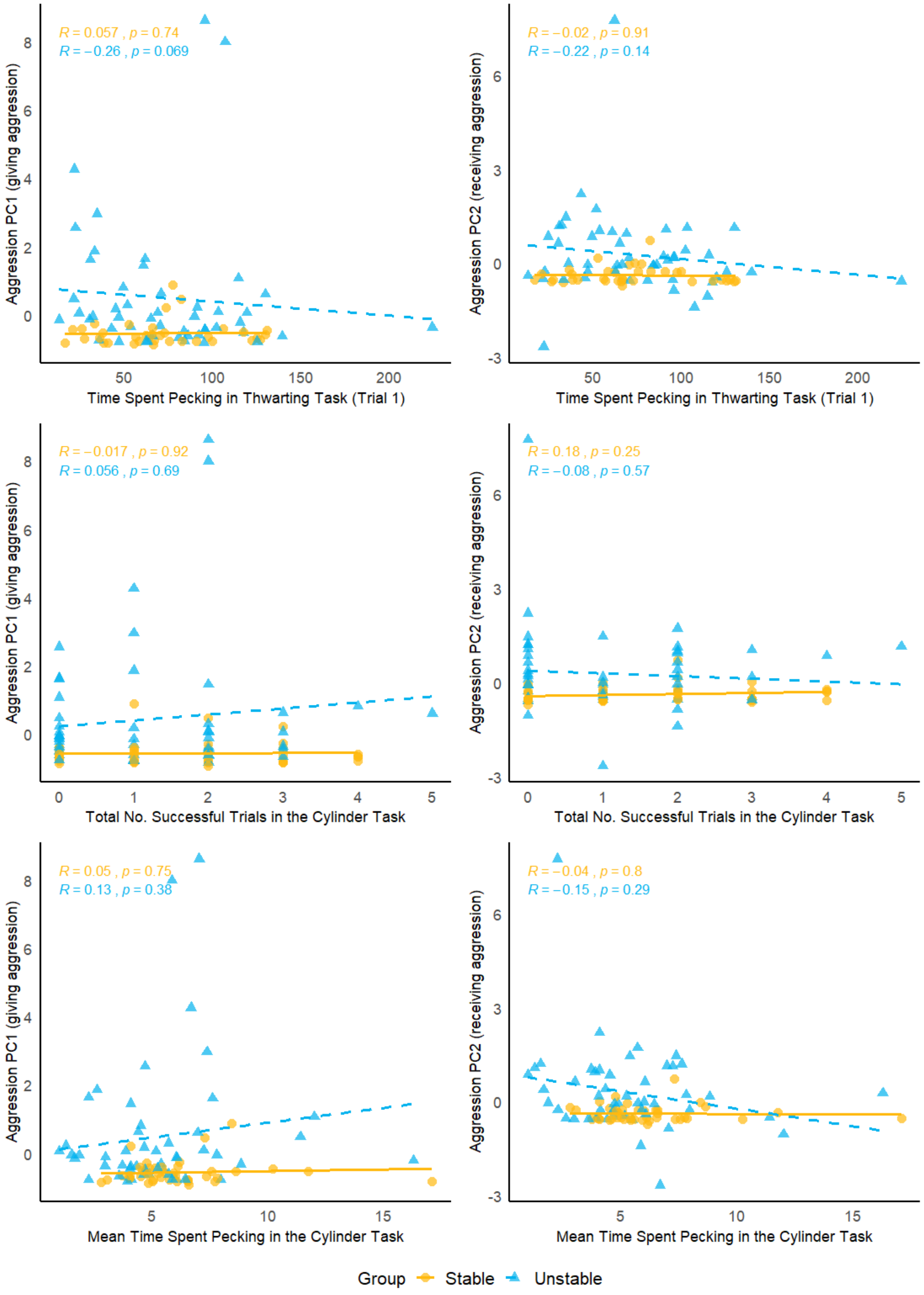


**Fig. S3** Correlations between aggression scores (left: PC1 – giving aggression; right: PC2 – receiving aggression) and response inhibition performance (top: time spent pecking in the thwarting task; middle: total successful trials in the cylinder task; bottom: mean time spent pecking in the cylinder task) for birds raised in stable and unstable groups, that were involved in aggressive interactions
